# Decomposing age effects in EEG alpha power

**DOI:** 10.1101/2021.05.26.445765

**Authors:** Marius Tröndle, Tzvetan Popov, Andreas Pedroni, Christian Pfeiffer, Zofia Barańczuk-Turska, Nicolas Langer

**Affiliations:** Department of Psychology, University of Zurich, Methods of Plasticity Research, Department of Psychology, University of Zurich, Zurich, Switzerland; University Research Priority Program (URPP) Dynamics of Healthy Aging, Zurich, Switzerland; Neuroscience Center Zurich (ZNZ), Zurich, Switzerland; Institute of Mathematics, University of Zurich, Switzerland

## Abstract

Increasing life expectancy is prompting the need to understand how the brain changes during healthy aging. Research utilizing Electroencephalography (EEG) has found that the power of alpha oscillations decrease from adulthood on. However, non-oscillatory (aperiodic) components in the data may confound results and thus require re-investigation of these findings. The present report aims at analyzing a pilot and two additional independent samples (total N = 533) of resting-state EEG from healthy young and elderly individuals. A newly developed algorithm will be utilized that allows the decomposition of the measured signal into aperiodic and aperiodic-adjusted signal components. By using multivariate sequential Bayesian updating of the age effect in each signal component, evidence across the datasets will be accumulated. It is hypothesized that previously reported age-related alpha power differences will disappear when absolute power is adjusted for the aperiodic signal component. Consequently, age-related differences in the intercept and slope of the aperiodic signal component are expected. Importantly, using a battery of neuropsychological tests, we will assess how the previously reported relationship between cognitive functions and alpha oscillations changes when taking the aperiodic signal into account; this will be done on data of the young and aged individuals separately. The aperiodic signal components and adjusted alpha parameters could potentially offer a promising biomarker for cognitive decline, thus finally the test–retest reliability of the aperiodic and aperiodic-adjusted signal components will be assessed.

## 1. Introduction

Alpha oscillations are by far the most widely studied phenomenon in human electrophysiology (EEG). Since the beginnings of EEG research in the 1920s, individual differences in alpha oscillations have been linked to variations in behavioral phenotypes and physiology (e.g., ref [1–3]). The alpha band is commonly defined as oscillatory activity in the range of frequencies between 8 and 13 Hz (e.g., ref [4,5]). These oscillations are typically observed over parietal and occipital electrode sites (e.g., ref [6,7]) and show the highest test—retest reliability of all frequency bands (e.g., ref [8]: *r* = 0.72-0.80). A robust finding is that alpha power decreases during task engagement (e.g., alpha event-related desynchronization). This led to the classical view that alpha amplitude reflects the idle state of cortical areas (for reviews see refs [1,3]). Moreover, modulations of alpha power have been linked to important concepts such as inhibition ([9], for reviews see refs [3,4,10,10]), attention [3,11], and memory retrieval [12].

Alpha oscillations are of particular interest in aging research because studies have repeatedly demonstrated that the alpha rhythm’s frequency slows (for a review, see ref [2]) and the alpha band power’s amplitude decreases with age [6,13–17]. A study investigating alpha power on source rather than on scalp level found age-related decreases particularly in posterior and occipital brain regions [13]. Furthermore, aging studies have often divided the alpha band into lower alpha and upper alpha sub-bands because these sub-bands have been linked to distinct cognitive functions [2,5,14,16,18]. Whereas the lower alpha band has been associated with attentional processing [18], age effects are found particularly in the upper alpha band and interpreted as changes of memory operations [14,16].

However, these results and interpretations of age-related alpha power changes need to be evaluated critically. Previous findings have demonstrated a slowing of the alpha rhythm with increasing age (e.g., ref [2]), so an age-related shift of the alpha peak frequency could lead to a reduction of alpha power as measured by fixed frequency bands. This may happen when the individual frequency center nears the lower limit of the fixed frequency band and therefore no longer captures the individual alpha oscillation (for a graphical illustration, see also Fig. 1 in [19]). Thus, the frequency band definition should be based on the individual alpha frequency center to rule out this confound.

**Figure 1:**
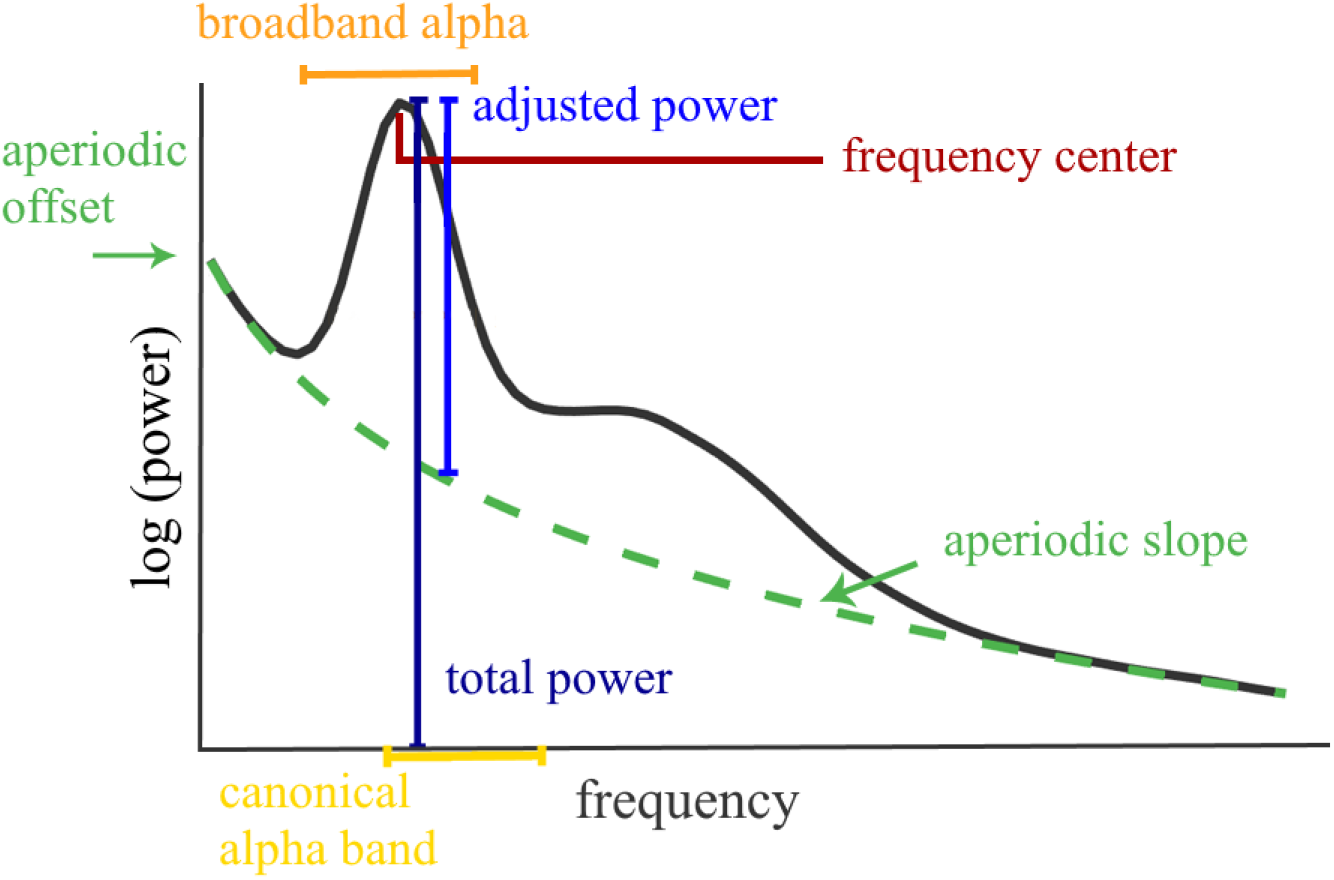
Illustration of a neural power spectrum (black line) and the various parameters extracted by the FOOOF algorithm. The yellow indicated canonical alpha band illustrates a fixed frequency band (e.g., the here used 8-13 Hz), the orange broadband alpha illustrates the here used band based on the IAF (−4Hz to +2Hz), which better captures the individual alpha oscillation.

More importantly, recent development in EEG signal processing techniques further question earlier findings of decreased alpha power in age [19]. It was demonstrated that the band power of the observed frequency consists not only of a periodic, or oscillatory, component but also of an aperiodic, or non-oscillatory, background signal. Therefore, band power should be investigated relative to this aperiodic signal [19–24]. The aperiodic signal is characterized by its shape (1/*f*), as its amplitude decreases with higher frequencies *f* (see Figure 1). This is based on the observation that EEG power spectra exhibit a static increase in power towards lower frequencies that has been shown to follow an underlying broadband power law with a negative slope [25]. In recent years, the aperiodic signal has attracted increasing attention from the research community. Whereas the offset of the aperiodic signal has been linked to general spiking activity (e.g., ref [26]) and to the blood-oxygen-level-dependent signal in functional magnetic resonance imaging [27] (fMRI), the slope has been associated with the synchronicity of activity in the underlying neural population [25,28]. A more asynchronous activation pattern has been shown to yield a flatter aperiodic slope. Furthermore, the aperiodic slope may be linked to the excitation–inhibition balance of transmembrane currents [29]. AMPA receptor mediated excitatory currents show high power at lower frequencies with a fast decay towards higher frequencies, leading to a steeper slope. GABA receptor mediated inhibitory currents show a slower decay of power towards higher frequencies, yielding a flatter slope for the aperiodic signal.

Thus, decomposing aperiodic activity from periodic appears mandatory in aging research as the aperiodic signal itself might change with increasing age [23]; hence, it may change the shape of the neural power spectrum even though the oscillatory pattern remains stable. Conventional analyses are prone to conflating periodic and aperiodic signal components and thus may lead to fallacious conclusions (see ref [19] for a discussion on further potential misinterpretations) and neurophysiological interpretations about age-related changes of alpha oscillations.

In this study we address this problem by directly comparing previously established age differences in unadjusted alpha power, here referred to as *total alpha power,* to age effects in the true periodic component of the alpha band, here referred to as *aperiodic-adjusted alpha power* and the aperiodic signal components, here referred as *aperiodic intercept and aperiodic slope* (see Table 2 for an overview of all terms related to the extracted alpha parameters).

So far, only one study compared age effects in the decomposed EEG signal using a small sample of 16 younger and 14 elderly subjects [19]. While a significant age-related decrease in aperiodic signal components, total and aperiodic-adjusted alpha power was reported; no statistical test was applied to investigate the particular contrast of age effects on total alpha power with age effects on aperiodic-adjusted alpha power. Thus, it remains largely unknown whether the age effects found on alpha power are mainly driven by aperiodic, periodic or both signal components. A rigorous statistical evaluation of age effects in alpha power, taking the aperiodic signal into account within a reasonably powered sample that allows to draw reliable and robust conclusions about these specific age differences, awaits demonstration.

The present report will fill this gap by evaluating a large sample of 100 young and 100 elderly subjects and employ multivariate statistical models to compare age differences in parieto-occipital total alpha power, aperiodic-adjusted alpha power, and aperiodic signal components. The multivariate approach is able to account for the potential correlation between the total and aperiodic-adjusted power measures. Total alpha power will be extracted using conventional spectral analysis, which does not adjust for the aperiodic background signal. Aperiodic-adjusted alpha power and aperiodic background signal components will be extracted using the Fitting Oscillations One Over F [19] (FOOOF) algorithm. Figure 1 illustrates the resulting parametrization of an EEG power spectrum. Alpha power measures will be extracted from a canonically defined fixed frequency band (8–13Hz) and from the individual anchor point (the individual alpha frequency IAF, see Table 2 for more details) to rule out confounds from a slowing IAF on the observed alpha power. Spatial patterns for each parameter will be examined by including all electrodes on the full scalp. Source analysis will enable the exploration of the neural generators of the aperiodic and aperiodic adjusted parameters. Results will be validated using a second, openly available dataset derived from 153 young and 74 elderly participants [30].

To qualify as biomarker of cognitive decline, a second key aspect is the reliability of the aperiodic signal or aperiodic-adjusted periodic activity, which remains unaddressed at present. Reliability estimates represent the ratio of within-subject variance (i.e., the measurement error) to the between-subject variance (i.e., difference between age groups). If the within-subject variance explains a large proportion of the total observed variance, drawing any conclusion about between-subject differences is precluded [31]. Neglecting reliability measures can thus result in costly studies that are unable to produce informative outcomes [31]. Therefore, it is a fundamental requirement to estimate reliability for these newly emerging measures.

In the current study, the test–retest reliability of the aperiodic intercept and slope and of the total and aperiodic-adjusted posterior alpha power will be assessed. The sample of 100 young and 100 elderly individuals will be used, from whom data was acquired at two consecutive measurements separated by a week.

Finally, the association between the decomposed power spectrum and cognitive functions will be examined. Literature has established a link between age-related decline in resting total alpha power with diminished attention and working memory performance [2,13,14,16,32,33]. However, these well-established findings were challenged by demonstrating that age-related cognitive decline is linked to a flattening of the aperiodic slope [23]. Thus, it remains unclear to what extent this relationship between alpha power and cognitive performance is in fact confounded by the aperiodic slope. To address this, neuropsychological tests assessing attention and working memory performance will be conducted in this sample of 100 young and 100 elderly participants. Subsequently, relationships to different measures of alpha power and the aperiodic signal will be examined.

The analysis code for the present study was implemented in a pilot dataset [34] before either of the two larger, independent datasets will be accessed or analyzed. All relevant analysis parameters will be fixed before conducting the main study, so there will be no degrees of freedom in the planned analyses, and overfitting errors will be minimized. Sequential Bayesian updating will be employed by fitting a Bayesian regression model to each of the three datasets and passing posterior distributions of each analysis to the next analysis as priors. Thus, evidence can be accumulated across the datasets. This will produce greater statistical power than independent analyses and more robust outcome parameters.

Based on previous literature and pilot data results, the present hypotheses are as follows:

- H1a: The alpha rhythm is slower (i.e., has lower alpha peak frequency) in the elderly group than in the young group.
- H1b: In the canonical, lower and upper alpha band, there is lower total power in the elderly group than in the young group.

The age differences in alpha power change when adjusting for the aperiodic signal:

- H2a: The upper and canonical alpha bands exhibit less age difference in aperiodic-adjusted alpha power than in total alpha power.
- H2b: The lower alpha band exhibits greater age difference in aperiodic-adjusted alpha power than in total alpha power.

Age differences in the aperiodic parameters are expected as follows:

- H3a: The aperiodic intercept is lower in the elderly group than in the young group.
- H3b: There is a flatter aperiodic signal (i.e., a smaller aperiodic slope) in the elderly group than in the young group.

With respect to the test-retest reliability:

- H4a: Both age groups exhibit a good to excellent test—retest reliability (i.e., intraclass correlation coefficient, ICC, > 0.6) for the total and aperiodic-adjusted alpha parameters and for the aperiodic slope and intercept.
- H4b: There are equivalent levels of test—retest reliability in the total and aperiodic-adjusted alpha parameters and in the aperiodic slope and intercept in both age groups.

With respect to the relationship between cognitive scores and the different parieto-occipital alpha power and aperiodic signal measures:

- H5a: Total alpha power is positively related to attention and working memory performance.
- H5b: This relationship is weaker when applied to aperiodic adjusted alpha power.
- H5c: The aperiodic signal slope is positively related to attention and working memory performance.

Our findings would have important implications for the interpretation of the underlying neurophysiological mechanisms of aging.

If we can demonstrate that the previously reported age-related decrease in alpha power is in fact confounded by alterations in the aperiodic signal (i.e., a decreased aperiodic intercept and a flattening of the aperiodic slope), as hypothesized in our present study, these results would challenge current neurophysiological interpretations of age-related decreases in alpha power (e.g., refs [13,35–38]). These interpretations link age-related decline in resting state alpha power to an increased excitability of thalamo-cortical and cortico-cortical interactions [39,40]. This increased excitability can be explained by the age-related gradual loss of cholinergic function in the basal forebrain (e.g., refs [41,42]). Importantly, experimentally impairing cholinergic forebrain function in animal models led to decreases in alpha power (e.g., refs [43,44]). In humans, this is consistent with observed decreases of alpha power in patients suffering from Alzheimer dementia and mild cognitive impairment, which are conditions that are characterized by impaired cholinergic forebrain function (e.g., refs [45–49]). The basal forebrain forms main cholinergic inputs to thalamic nuclei [50], which are considered key structures in the generation of cortical alpha oscillations (e.g., refs [51,52]). Thus, it is hypothesized that the decreased cholinergic input leads to decreases of power in the cortical alpha oscillations. This hypothesis is supported by work in animal models, which demonstrates that stimulation of cholinergic receptors in the reticular thalamic nucleus and thalamo-cortical cells produces alpha oscillatory activity [53].

If the decomposition of the measured EEG signal in the present study will reveal that age-related differences only relate to the aperiodic signal component but diminish for the periodic signal component (i.e., adjusted alpha power), as hypothesized in our present study, the current cholinergic theory on the effects of aging on the EEG signal needs to be reconsidered.

An age-related decrease in the aperiodic signal intercept would indicate that age differences in the measured EEG signal are rather caused by decreased neuronal population spiking activity [19,54]. Decreases in the aperiodic slope can be summarized in the neural noise hypothesis of aging [26]. This hypothesis states that age-related cognitive decline is caused by increased occurrence of temporally decorrelated spikes (i.e., noise) which are not involved in encoding of information or network communication. These irregular spikes decouple the synchronization mechanisms between brain regions, which leads to communication errors. This increased occurrence of errors in the communication between brain regions in turn result in age-related cognitive deficits [26]. The irregular spikes are caused by greater local positive excitatory feedback, driven by an increased excitation-inhibition ratio [26]. An increased neural excitation-inhibition ratio has been linked to a flattened aperiodic slope both in computational models and animal studies [29]. In line with this theory are findings that show the decrease of the aperiodic slope to be associated with more asynchronous activation patterns in neural populations [25,28]. If the present study will demonstrate that resting alpha oscillatory power associations with cognitive scores are also conflated with effects in the aperiodic signal, it would further support the theory that increased neural noise is the driving physiological mechanism underlying age-related cognitive decline.

Taken together, preserved alpha power together with age differences in the aperiodic signal component would indicate that previous interpretations of age difference in the measured EEG signal are rather caused by different, so far neglected physiological mechanisms. Thus, age-related changes in neural noise need to be incorporated into the above described existing interpretations.

### 1.1 Summary table

**Table 1.**
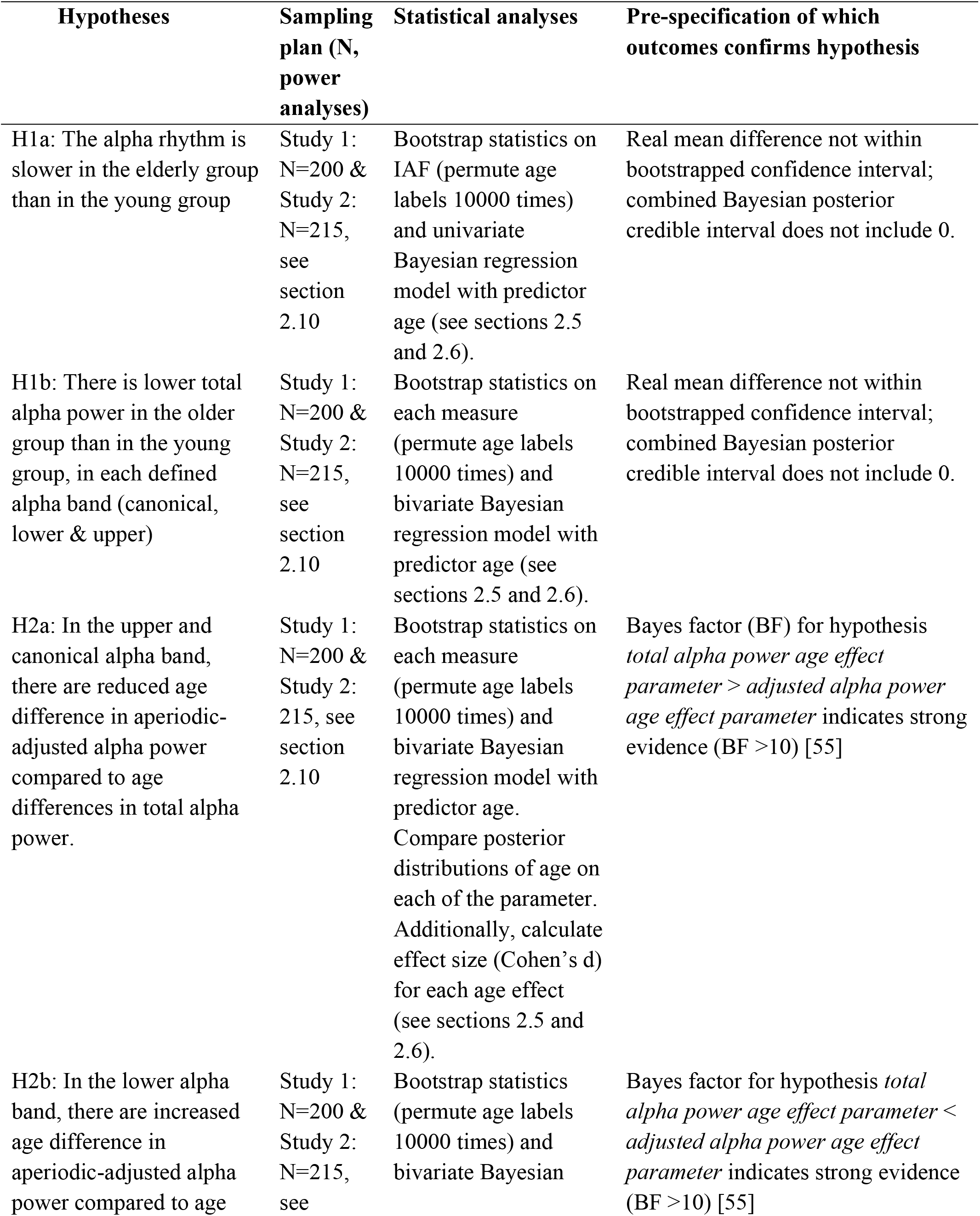
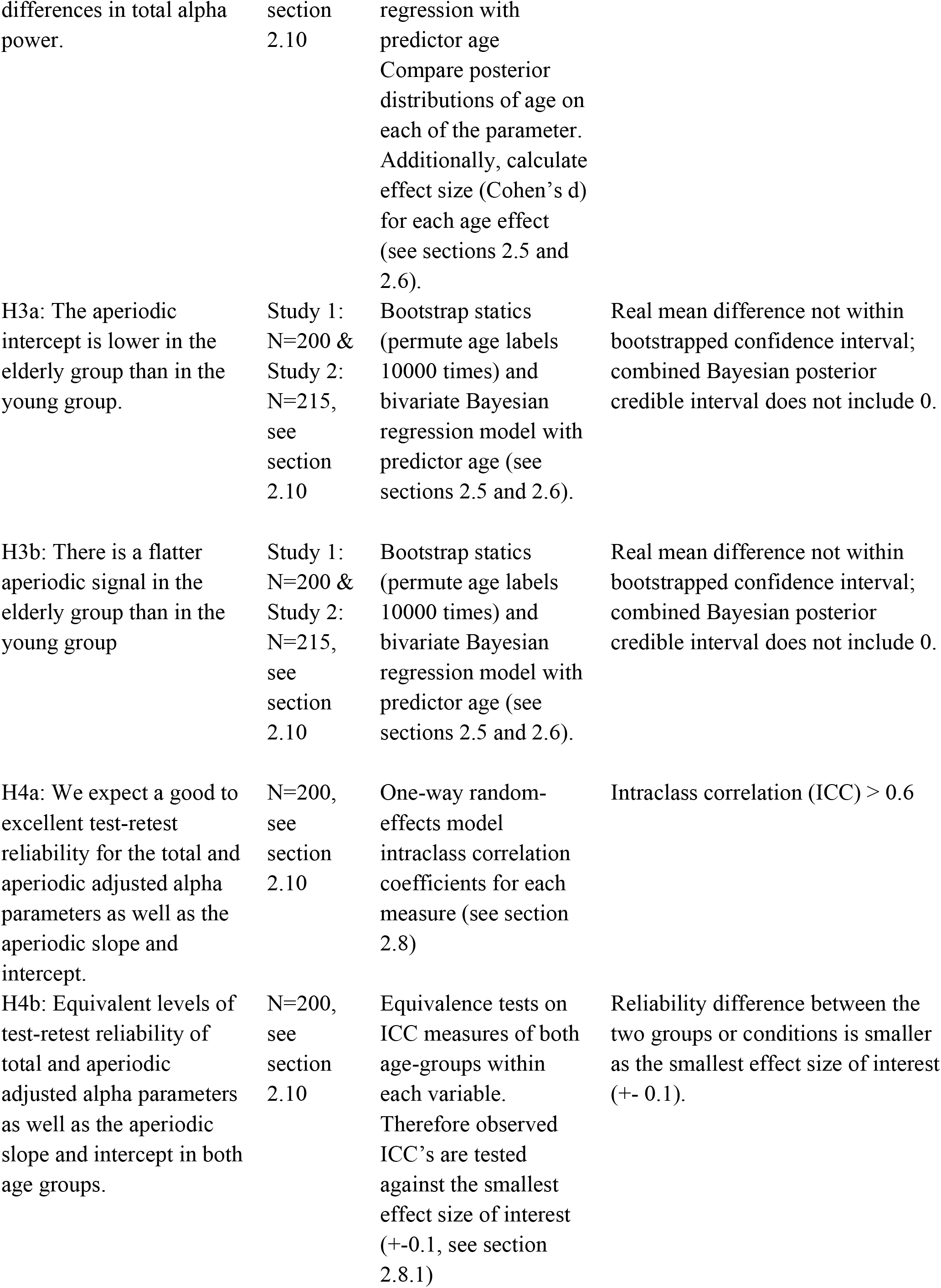
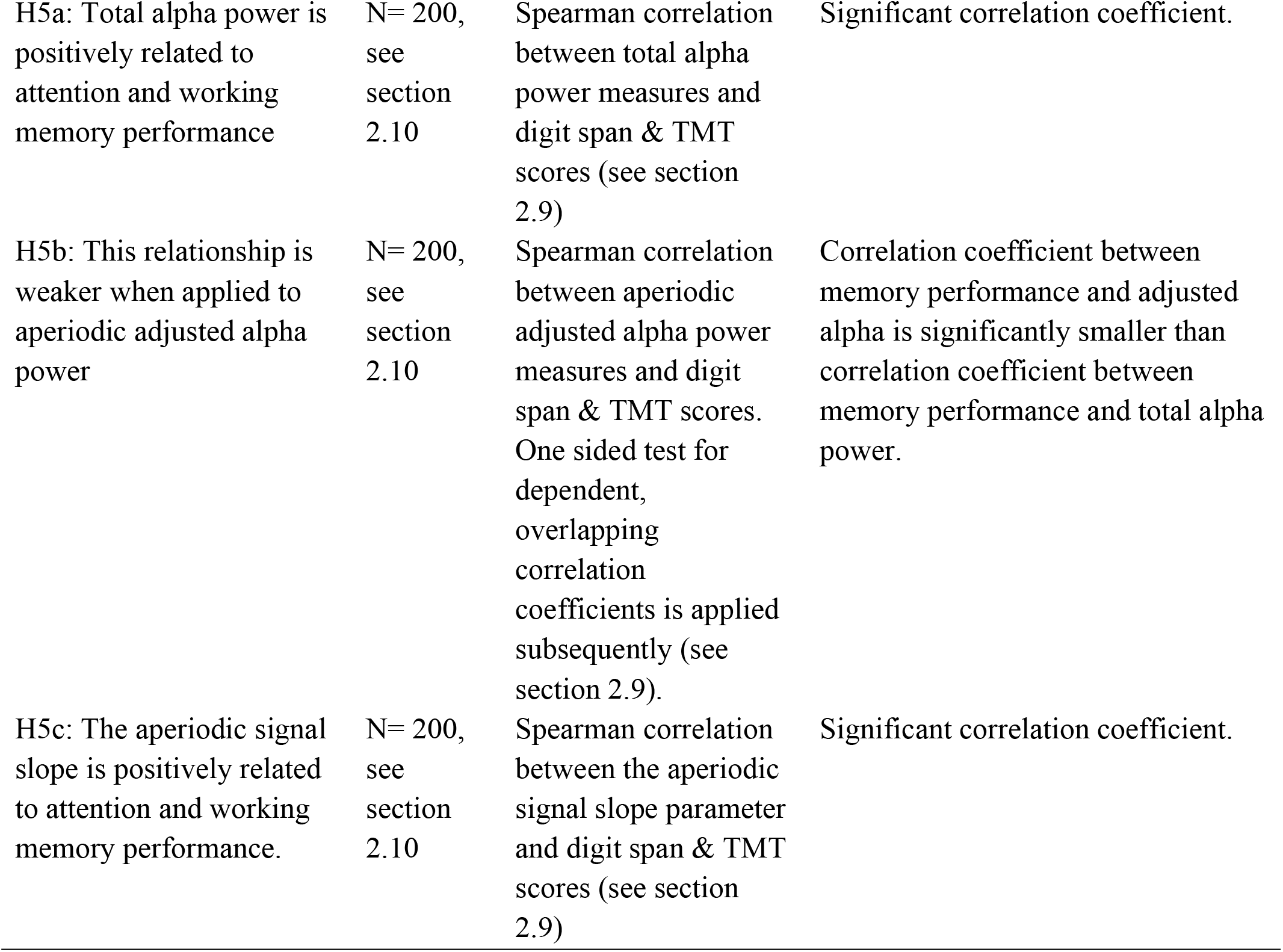
Summary of each hypothesis, the according sampling plan, proposed statistical analyses and a pre-specification of which outcome will confirm or disconfirm the specific hypothesis.

## 2. Methods

### 2.1. Datasets

This study will use data that is currently being recorded in our laboratory as part of a larger project. The larger project aims to quantify healthy-aging-related task performance and neuroelectric correlates in seven EEG tasks as well as in EEG resting-state recordings. The tasks aim to measure working memory, processing speed, and inhibitory control. One hundred healthy elderly subjects (60–80 years) and 100 healthy young participants (18–35 years) will be recruited. Exclusion criteria are suffering from psychiatric symptoms, severe neurological disorders such as epilepsy, prior head injuries, a stroke, a transient circulatory disorder of the brain, diagnosis of dementia, Huntington’s disease, Parkinson’s disease, current use of psychotropic drugs such as antidepressants, alpha-agonists, neuroleptics, and mood stabilizers, and intake of any recreational drugs. Additionally, subjects whose score in the mini-mental state examination [56] (MMSE) is below 27 will be excluded due to the high risk of dementia or mild cognitive impairment [57]. Each subject will take part in two experimental sessions following the identical EEG protocol. This is done to assess the test–retest reliability of all dependent measures. The sessions are scheduled with an inter-session interval of one to two weeks and at the same time of the day. The local ethics committee has approved the project, and all participants will give their written informed consent to participate in the study. The present study will analyze resting-state EEG data of participants who take part in both recording sessions: 400 recordings of 200 participants.

Additionally, a second independent dataset will serve the validation of the results obtained in our laboratory. This openly available, published dataset [30] contains resting-state EEG measurements of 215 healthy participants comprising a young and an elderly group (*N*_young_ = 153, mean age = 25.1 years, sd = 3.1, age range = 20–35 years, 45 female; *N*_eldery_ = 74, mean age = 67.6 years, sd = 4.7, age range = 59–77 years, 37 female). These two datasets have not been evaluated in any way and will be analyzed only after in principle acceptance of the present manuscript.

A third dataset was used for the pilot analysis (see below). This dataset originates from a previous study [34] and contains EEG recordings of 118 subjects (*N*_young_= 63, mean age = 23.37 years, sd = 3.91, age range = 18–35 years, 40 female; *N*_old_ = 55, mean age = 68.40 years, sd = 3.29, age range = 61–77 years, 23 female). More details about the pilot dataset are provided below (see section 3.1).

### 2.2. Experimental setup and procedure

Prior to the EEG assessment, participants perform a cognitive test battery. During EEG acquisition individuals are comfortably seated in a chair in a sound- and electrically shielded Faraday recording cage. The cage is equipped with a chinrest to minimize head movements and a 24-inch monitor displaying a fixation cross. Participants are informed that EEG is recorded while they rest with their eyes alternately open or closed. Instructions to open or close the eyes are automatically presented via cage intern loudspeakers. Adopting the recording protocol from previous studies [34,58], within five repetitions, participants are asked to fixate for 20 s followed by 40 s of eyes closed recordings. This results in 100 seconds eyes open and 200 seconds eyes closed data available for further analysis.

### 2.3. Cognitive assessment

The cognitive assessment will include the Trial making Test A / B (TMT, e.g., ref [59]) and digit span forward / backward task [60].

#### TMT A/B task

In the TMT A task, the participant is asked to increasingly connect a series of 25 spatially distributed circled numbers using a pencil. In the B part, 25 circled numbers and letters need to be connected alternatingly in a numerical and alphabetical order. Time to completion is measured by the experimenter. The subtest TMT A measures motor speed and visuospatial attention, the B version further captures working memory operations such as executive control (e.g., refs [61,62]). The score of the TMT A and B versions are defined as the time to completion in each condition. Additionally, the ratio of TMT B / TMT A will be calculated as a third score. This is thought to minimize visual attention components and to be a more direct indicator of working memory function [63].

#### Digit span task

The digit span task constitutes a forward and backwards version. A sequence of numbers is read to the participant who is then asked to repeat the sequence either forward or in reverse order. The length of the sequence increases until the participant fails to recall the correct order twice (for more details, see ref [60]). The forward task is designed to measure attention and short term memory, or the functioning of the phonological loop component of Baddely’s working memory model [64]. The digit span backwards task captures working memory processes such as executive control functions (e.g., refs [65,66]). The performance score (forward and backward) will be defined as the number of correctly recalled sequences in each condition.

### 2.4. Electroencephalography acquisition and preprocessing

The high-density EEG is recorded at a sampling rate of 500 Hz, using a 128-channel EEG Geodesic Sensor Net system (Electrical Geodesics, Eugene, Oregon). The recording reference is at Cz (vertex of the head), and impedances are kept below 40 kΩ. All subsequent analyses will be performed using MATLAB 2018b (The MathWorks, Inc., Natick, Massachusetts, United States). EEG data will be automatically preprocessed using the current version (2.5) of the MATLAB toolbox Automagic [67]. The analysis pipeline will consist of the following steps. First, error-prone channels will be detected by the algorithms implemented in the eeglab plugin clean_rawdata (http://sccn.ucsd.edu/wiki/Plugin_list_process). An electrode is defined as an error-prone when recorded data from that electrode is correlated at less than 0.85 to an estimate based on neighboring electrodes. Furthermore, an electrode is defined as error-prone if it has more line noise relative to its signal than all other electrodes (4 standard deviations). Finally, if an electrode has a longer flat line than 5 s, it is considered error prone. These error-prone electrodes will automatically be removed and later be interpolated using a spherical spline interpolation (EEGLAB function eeg_interp.m). This interpolation will be performed as a final step before the automatic quality assessment of the EEG files (see below). Next, data will be filtered using a high-pass filter (cutoff frequency (−6 dB): 0.5 Hz) and Zapline [68] will be applied to remove line noise artifacts, removing 7 power line components. Subsequently, independent component analysis (ICA) will be performed on temporary highpass filtered data (cutoff frequency (−6 dB): 1 Hz). Components reflecting artifactual activity will be classified by the pre-trained classifier ICLabel [69]. Each component being classified with a probability rating >0.8 for any class of artifacts (line noise, channel noise, muscle activity, eye activity or heart artifacts) will be removed from the data. The retained components will be back-projected on the 0.5 Hz high-pass filtered data. Finally, residual bad electrodes will be excluded if their standard deviation exceeds a threshold of 25 μV. After this, the pipeline automatically assesses the quality of the resulting EEG files based on four criteria. First, a data file will be marked as bad-quality EEG and will not be included in the analysis if the proportion of high-amplitude data points in the signals (>30 μV) is larger than 0.2. Second, more than 20% of time points show a variance larger than 15 μV across channels. Third, 30% of the electrodes show high variance (>15 μV). Fourth, the ratio of bad electrodes is higher than 0.3. This standardized and objective preprocessing pipeline and data quality metrics remove all degrees of freedom from the preprocessing. The analysis code for the planned preprocessing can be found in an OSF repository (https://osf.io/8e2kd/) and will be applied on all three investigated datasets. After this automatic preprocessing step, the resulting continuous EEG data will be down-sampled to 125 Hz, and the number of electrode will be reduced to a subset of the 70 electrodes that closely match the standard 10-10 electrode locations [70]. This will be done to reduce computational costs for parametrization of the neural power spectra and estimation of the Bayesian statistical models. It will also allow direct comparison of EEG from the three datasets using different cap layouts: ours described above, that of the validation dataset [30] and that of the pilot dataset [34]. Finally, the data will be re-referenced to a common average reference. The first and the last 2 s of each eyes-closed block will be discarded to exclude motor activity related to opening and closing the eyes and auditory activity due to the prompt from the loudspeakers. The remaining data will be concatenated, resulting in a total of 180 seconds of continuous EEG data. Subsequently, the data is further segmented into epochs of 2 sec length. EEG epochs exceeding a ±90μV amplitude threshold will be excluded from further analysis. We note that, because several quality metrics from the Automagic toolbox were used for selecting EEG data, it is unlikely that at this point a significant number of EEG epochs will be excluded. Nevertheless, we will exclude subjects from the analysis for whom less than 50% of the epochs for the eyes closed condition remain available for further analyses.

### 2.5. Spectral analysis

Spectral analysis will be performed on artifact-free 2 sec segments from five blocks of the eyes-closed condition. Only data from the eyes closed condition will be analyzed, as this data contains fewer artifacts and generally shows the strongest alpha amplitudes. Additionally, eyes-closed data has been previously used to establish a link between age and alpha power [6,13,14,17]. Power spectral densities (PSDs) will then be calculated using Welch’s Method [71] as implemented in the EEGLab toolbox [72] (by default, non-overlapping windows). Zero padding will be applied to provide a frequency resolution of 0.25 Hz in the 2 s time windows. Averaging the individual PSDs of each window results in a smoothed power spectrum that complies with the requirements of the FOOOF algorithm used subsequently (see below, Figure 1). Additionally, PSDs will be transformed to log scale, to make results equally scaled to outputs from the FOOOF algorithm, which only operates in log space. In the following, the two approaches to extract total alpha power and the adjusted alpha power together with the aperiodic signal are described. Table 2 provides an overview of all extracted parameters.

**Table 2.** Overview of extracted parameters.

| Parameter of uncorrected (“total”) PSD | Parameter from aperiodic adjusted PSD | Description |
| --- | --- | --- |
| Total canonical alpha power | Aperiodic adjusted canonical alpha power | Averaged log power in 8-13Hz window |
| Total lower alpha power | Aperiodic adjusted lower alpha power | Averaged log power in window [-4 Hz IAF] |
| Total upper alpha power | Aperiodic adjusted upper alpha power | Averaged log power in window [ IAF +2Hz] |
| Individual alpha frequency | Individual alpha frequency | Frequency at maximum power in search window 7-14Hz |
| N/A | Aperiodic intercept | Intercept parameter of the aperiodic signal component |
| N/A | Aperiodic slope | Slope parameter of the aperiodic signal component |

*Note:* Individual alpha frequency is not affected by the adjustment for the aperiodic signal. If the search criteria in the total power spectrum cannot be met, IAF will also not be taken from the flattened (aperiodic adjusted) power spectrum. Aperiodic intercept and slope are not applicable on the uncorrected PSD.

#### 2.5.1. Computation of total alpha power

In this standard analysis approach, no adjustment will be made for the aperiodic signal, as it was used throughout previous literature. First, the IAF will be determined by extracting the maximum power value in between a lower and upper frequency limit [2]. Following previous work, these frequencies limits will be set to 7 and 14 Hz [73,74]. If this maximum is at the border of the search range, no IAF will be extracted for that subject and the corresponding data will be excluded from further analysis (see also exclusion criteria in section 2.5.4). Therefore, the effective search range for the IAF will be between 7.5 and 13.5 Hz. If an IAF can be identified, additional alpha sub-bands will be extracted: *total lower alpha power* [−4 Hz IAF] and *total upper alpha power* [IAF +2 Hz] will be calculated by averaging power in the above-defined range in reference to the IAF [2]. Additionally, the commonly used *total canonical alpha band power* will be calculated by averaging power in the range fixed of 8 to 13 Hz [13].

#### 2.5.2. FOOOF algorithm

The FOOOF algorithm [19] parameterizes the power spectrum to separate oscillations from the aperiodic background signal. The algorithm estimates oscillatory peaks that are superimposed on the aperiodic background signal (see Figure 1) and therefore measured relative to this rather than to the absolute zero. Thus, it parametrizes the PSD by iteratively fitting the aperiodic background curve (L) to the observed smoothed spectral signal, resulting in two parameters: the aperiodic intercept b and the aperiodic exponent χ (i.e., slope, the smaller χ, the flatter the spectrum).

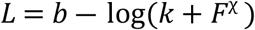

Here, F represents the vector of input frequencies and k the “knee” parameter, which is not further discussed here, as it is set to 0 in the proposed analysis (i.e., no bend of the aperiodic component is additionally modeled in the data, which is the default state of the FOOOF algorithm).

In order to extract oscillatory components, this aperiodic background signal is subtracted from the power spectrum. Gaussians are iteratively fitted to the remaining signal and subsequently subtracted whenever data points exceed two standard deviations of the data. The Gaussians represent the true oscillatory components in the data; if data points are below the specified threshold, they are considered as noise. This results in a data-driven number of Gaussians, each parameterized by the frequency center, power relative to the aperiodic signal and the frequency bandwidth. The power spectrum is therefore modeled by

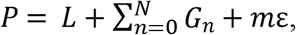

where *G*_n_ represents the *n*^th^ Gaussian, and *m* the scaling factor of the noise ε. Note that this description of the algorithm is simplified; for a more detailed definition, see ref [19].

In the planned analyses, the frequency range of 2 to 40 Hz will be passed to the algorithm because very low frequencies may lead to overfitting of noise as small bandwidth peaks. The release 1.0.0 of the FOOOF toolbox from the github repository (https://github.com/fooof-tools/fooof) will be used, applying standard peak detection parameters (2 standard deviations above mean).

To correct the above-described total alpha power measures (see 2.5.1) for the aperiodic signal, the aperiodic signal will be reconstructed by its parameters. Subsequently, the aperiodic signal will be subtracted from the total power spectrum to receive an aperiodic adjusted power spectrum. Using this power spectrum, adjusted alpha power values *(adjusted canonical alpha power, adjusted lower alpha power, adjusted upper alpha power)* will be calculated in the frequency ranges described above (2.5.1).

#### 2.5.3. Cluster-wise analysis

To test age effects in electrode sites derived from literature, electrode-cluster-based analyses will be performed. This cluster will be based on data from the parietal and occipital electrodes: E72 (POz), E75 (Oz), E62 (Pz), E67 (PO3), E77 (PO4), here referred to as the parieto-occipital cluster (see Figure 3a). These electrodes were chosen because of the strong prominence of Oz and Pz electrodes in EEG alpha peak research [2] and previous findings for age effects on alpha band power in these electrodes (e.g., refs [6,14]). To account for individual anatomical differences and to create a more robust cluster, the three electrodes adjacent to Oz (E75) and Pz (E62) will be added (E72 (POz), E67 (PO3), E77 (PO4)). All the parameters described above will be averaged within the cluster.

#### 2.5.4. Exclusion criteria

Before statistical analyses are performed, data will be excluded if any the following criteria applies:

- The fit of the parameterized power spectrum to the original PSD is below a cut-off of R^2^ < 0.9. If the fit of the parametrized spectrum is below the specified cut off, total alpha parameters will be still included, as these are not contingent on the parametrization of the FOOOF algorithm.
- Any of the extracted parameters exceed a threshold of 3 standard deviations above or below the mean of the sample.
- No individual alpha peak can be detected. If no alpha peak can be detected, aperiodic signal components of the according subject will still be used, as the aperiodic signal components do not depend on the individual alpha peak detection.
- EEG data file is rated “bad” by the preprocessing pipeline.
- Less than 50% artifact-free data.

These first three criteria will be applied within data of each electrode separately.

### 2.6 Statistical analyses of age differences

To correct for multiple comparisons in the statistical analyses, the significance level will be adjusted. We assume a high correlation between the eleven outcome variables, as many of the dependent variables represent different characteristics of the individual alpha oscillations. To account for this, we will first calculate the effective number of tests of all dependent variables using Nyholt’s approach [75]. Following this approach, the significance level (0.05) will then be adjusted using Šidák-Correction [75]. Subsequently, the confidence intervals of the bootstrap statistics (see 2.6.1) as well as the credible intervals (CIs) of the Bayesian posterior distributions (see 2.6.2) will be defined based on the newly calculated levels of significance.

Besides the electrode cluster based analysis, we will also investigate the spatial distribution of all statistical parameters on full scalp level. To correct for multiple comparisons we will apply a nonparametric cluster-based permutation analysis. This will be done using the ft_freqstatistics function implemented in FieldTrip [76].

#### 2.6.1 Bootstrap analysis

For each dataset, robust and assumption free bootstrap statistics will be performed. Therefore the original data will be permuted 10000 times. To investigate age effects within each of the nine above-described parameters (see Table 2), the mean age difference of all nine parameters will be calculated within each permuted dataset. Subsequently, the corrected confidence interval of the bootstrapped age differences will be calculated for each measure. If this interval does not include zero, the age difference will be considered significant.

#### 2.6.2 Bayesian regression model

In order to accumulate evidence across the three different dataset used in this study, an additional multivariate Bayesian generalized linear mixed model was formulated using the brms R package [77]. In this analysis, the model will first be fitted to the pilot data, using uninformative priors (see below). The extracted posteriors will not be interpreted but used as priors for the next analysis of the main dataset (N=200), which uses the same model as before. This will be done by applying the best fitting distribution to the posterior samples using the fitdistrplus R package [78]. The resulting posterior distributions of the main analyses will be approximated the same way and then serve as priors for the analyses of the validation dataset [30] (N=215). Only these resulting posterior distributions of the age effects will be interpreted as a final and more robust outcome.

The multivariate model was chosen as it is able to account for correlation between multiple dependent variables. Further advantages of this Bayesian approach are the facilitations to make inferences about the nonexistence of any effects and to statistically compare posterior distributions of the different parameters. In all planned analyses, the following model will be used. To account for the repeated measurement structure of the study design of the main analysis (two identical measurements of each subjects within 1-2 weeks) and the multiple dependent variables per subject, random intercepts are added for the participant IDs as shown in equation 1:

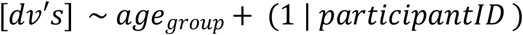

This model will be fitted five times in each of the three datasets: for each of the three alpha band power parameters together with their according aperiodic adjusted equivalent (lower alpha, upper alpha, canonical alpha), for the aperiodic signal components together (intercept and slope) and for the IAF. This will allow the direct comparison between the resulting posteriors of each total alpha parameter and its corresponding aperiodic adjusted equivalent using the brms hypothesis function (see below).

If the corrected posterior CI of a parameter of the model will not include 0, it will be considered significant. This will allow to test hypotheses H1a, H1b, H3a and H3b. If the CI will include zero, the test for practical equivalence [55,79] will be used to assess whether the observed effect is in favor of the null hypothesis. The test for practical equivalence is based on the “HDI+ROPE” decision rule [55,79] to decide whether parameter values should be accepted or rejected against an explicitly formulated null hypothesis (i.e., region of practical equivalence “ROPE”). If the ROPE completely covers the 89% highest density interval (HDI, i.e., credible values of a parameter are inside the ROPE) the null hypothesis is accepted. According to Kruschke’s recommendation [79], the ROPE will be set to a negligible effect size (d = −0.1 to d = 0.1) [80]. Converting this effect size to the betas provided by the models (standard deviation of the dependent variable is 0.5, see below), yields a ROPE of – 0.05 to 0.05 [81].

Furthermore, to test whether age differences are changing when adjusting alpha parameters for the aperiodic signal (H2a & H2b), the according posterior distributions will be compared. Therefore, one sided Bayes Factors will be calculated for the hypotheses *total alpha power age effect parameter* > *adjusted alpha power age effect parameter* or *total alpha power age effect parameter* < *adjusted alpha power age effect parameter*, depending on the corresponding above-described hypotheses H2a and H2b. This will be done using the brms hypothesis function [77]. Additionally, to estimate how strongly the age differences change, Cohens d’s will be calculated and reported for the age differences within each alpha measure.

In line with Gelman’s recommendations [82], the predictors and outcome variables of the Bayesian regression model will be scaled as follows: The binary parameter (age) will be centered at 0 and each numeric parameter (total canonical alpha power, adjusted canonical alpha power, individual alpha frequency, aperiodic signal slope and aperiodic signal intercept) will be scaled to provide a mean of 0 and standard deviation 0.5. For the first analysis of the pilot data, weakly informative Cauchy priors (mean = 0, scale = 2.6.2) were chosen in line with Gelman’s recommendations [82] for Bayesian regression models.

### 2.7 Source level analysis

To investigate the neural generators of adjusted alpha power and aperiodic slope, source level analysis utilizing a beamformer spatial filtering approach will be applied. A template forward model will be derived from the MNI ICBM 2009 template brain (http://www.bic.mni.mcgill.ca/ServicesAtlases/ICBM152NLin2009) using the OpenMEEG implementation [83] of the Boundary Element Method (BEM). The linear constrained minimum variance [84] (LCMV) beamformer algorithm will be applied to construct the spatial filters. These spatial filters will then be multiplied with the eyes closed time series data, resulting in source level time series for each voxel. Subsequently, source power spectra will be calculated and decomposed into aperiodic and periodic signal components as described in 2.5. Based on these, aperiodic adjusted canonical alpha power and the aperiodic slope parameter will be visualized.

### 2.8 Test-retest reliability

In order to quantify test–retest reliability for the output measures collected at the two recording sessions per subject, we will calculate one-way random-effects model intraclass correlation coefficients (ICCs). This will be done using the absolute agreement measure among multiple observations [85–87] with the irr open-source software package (https://CRAN.R-project.org/package=irr). These measures will be calculated for the eleven parameters of the parietooccipital electrode cluster. We will use the generally adopted interpretation of ICC [88]: less than 0.40 (poor reliability), between 0.40 and 0.59 (fair reliability), between 0.60 and 0.74 (good reliability), and between 0.75 and 1.00 (excellent reliability).

Additionally, Bland-Altman plots [89] will be used for graphical comparison of the measurements from test and retest recording sessions. In the Bland-Altman plot, each sample is represented on the graph by plotting the mean value of the two assessments against the difference between them. The chart can then highlight possible anomalies, such as that one method overestimates high values and underestimates low value [90]. We will also use a quantitative method to assess the agreement of test and retest. This is based on a priori defined limits of agreement: as for other relevant measures, it is recommended that 95% of the data points should lie within ±1.96 SD of the mean difference [91,92].

#### 2.8.1 Equivalence test

To investigate the agreement between the reliability of the two age groups (hypothesis H4b: test-retest reliability between the two ages groups is not different) equivalence tests on the ICC values will be conducted [93]. The equivalence test seeks to determine whether the reliability difference between the two groups or conditions is at least as extreme as the smallest effect size of interest (SESOI), following the two one-sided tests (TOST) procedure. In other words, the equivalence test does not test whether there exists no reliability differences at all between the groups, rather does it examine whether the hypothesis that effects are extreme enough to be considered meaningful can be rejected [93]. Performing TOST therefore involves determining the SESOI. Thus, SESOI and its lower and higher equivalence bounds respectively need to be determined first. As ICC values can be roughly compared to Cohen’s d effect size measures, we will consider an ICC of +-0.1 as the respective equivalence bounds. The TOST procedure is then performed against lower and upper equivalence bounds that are specified based on the SESOI.

### 2.9 Relation to cognitive scores

To test hypotheses 5a, 5b and 5c, Spearman correlations between the different measures of parietooccipital alpha power and the aperiodic signal (total and aperiodic adjusted lower-, upper-, and canonical alpha power, aperiodic slope and intercept) and the scores of the TMT and the digit span task will be calculated. Spearman correlation is preferred over Pearson correlation to avoid effects of outliers and to minimize bias due to possible non-normality of the data. To test hypothesis 5b, we will test whether correlation coefficients between adjusted alpha power measures and the cognitive scores are smaller than correlation coefficients between total alpha power measures and the cognitive scores. Therefore, a confidence interval based one sided test for dependent, overlapping correlation coefficients implemented in the R cocor package [94] will be applied [95]. If there will be no significant difference, equivalence tests will additionally be performed as described in 2.8.1. SEOSI will be defined as r=0.1, which is the smallest correlation coefficient acknowledged as a meaningful effect by Cohen [96]. This procedure will be applied to data of the young and elderly group separately.

### 2.10 Power analysis

In order to determine statistical power in the available sample, a literature search was conducted. Subsequently a simulation-based power analysis was performed. The literature search focused on age differences in the following EEG features: spectral power in the alpha frequency range as well as parameters extracted by the FOOOF algorithm (i.e., aperiodic signal slope and intercept). Five studies were identified which reported analyses similar to the planned analyses on age differences in total alpha spectral power [6,13–15,17].

Ref [15] reported a *p*-value *(p* < 0.01) but no measure that could be used to calculate an effect size. Ref [6] reported *t*-values of interest as a topographical plot in which the color scale represented individual *t*-values. From this color scale, no exact values can be extracted for further calculation. Ref [17] provide unstandardized regression coefficients but these do not allow any calculation of an effect size. The remaining two studies [13,14] both divided the alpha band into a lower and an upper sub-band. Ref [14] reported a negative correlation of *r* = -.27 between upper alpha band power and age in a parietal electrode (Pz). Ref [13] calculated alpha power on source space. The upper alpha band was investigated in the occipital and parietal brain regions which most closely match the electrode cluster in the planned study. For the occipital cluster, an *r*^2^ of .18 (r = ± .42) was reported; for the parietal cluster, the corresponding *r*^2^ value was .1 (*r* = ± .32). Taken together, the literature review revealed five studies reporting consistent age differences in total alpha power, but only two allowed a calculation of effect sizes. Due to the small number of available observations and the high probability of publication bias (e.g., ref [97]), only the smallest observed effect (*r* = -.27) was chosen as the basis for power analysis. This value was transformed to Cohen’s *d,* resulting in *d* = 0.56.

The literature search for age differences in aperiodic background signal parameters extracted as in the FOOOF algorithm yielded only one study. Ref [23] reported a correlation of aperiodic signal slope with age of *r* = .66 (*d* = 1.75). Therefore, we conclude that the minimum sample size needed for the planned study is driven by the age differences in total alpha power (*d* = 0.56), as all other effects are larger and therefore require smaller sample sizes.

Next, a retrospective power analysis was performed, as the maximum number of participants available for the main study is 200. Therefore, a simulation-based approach was chosen. One thousand simulated datasets of alpha parameters were drawn randomly from normal distributions, each with a group difference of *d* = 0.56. On each dataset, a brms model was fitted investigating age effects on the alpha power. When the 95% CI of the model coefficient of the age effect did not include zero, the result was considered significant [55,98]. When using a sample size of 200, the age effect was found in 97.4% of the 1000 simulations. Figure 2 shows the results of the simulations using sample sizes ranging from 2 to 200.

**Figure 2:**
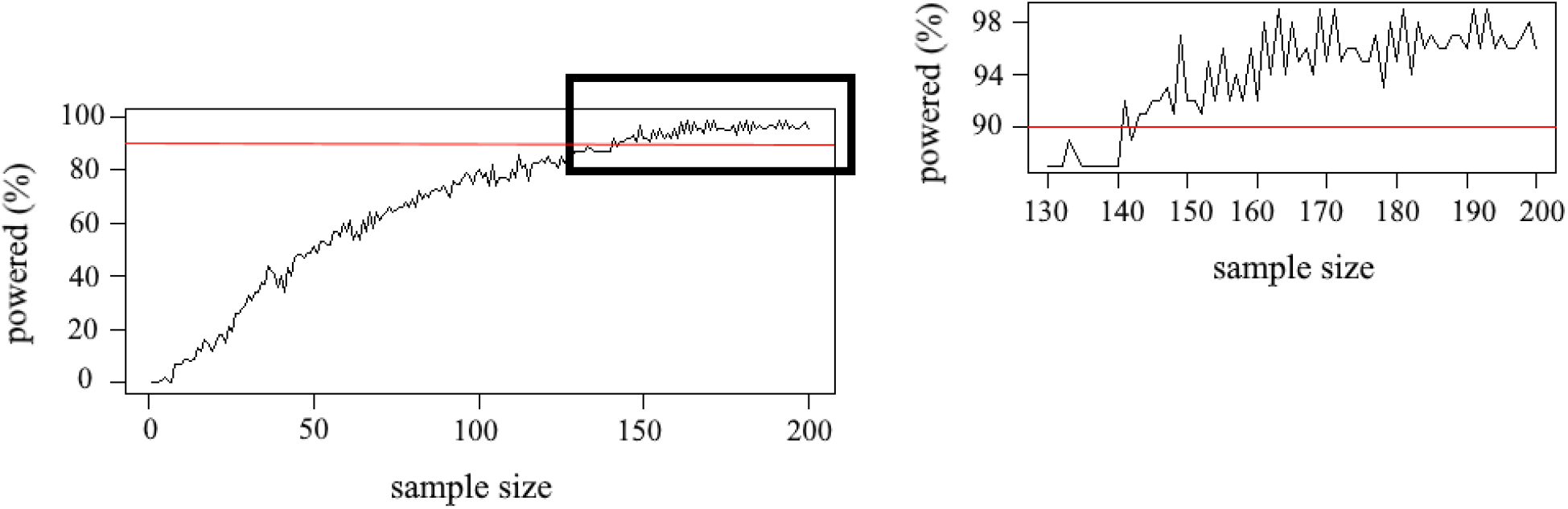
Percentage of simulations (1000 per sample size) being sufficiently powered (i.e., showing significant age effects on total alpha power). Red line indicates power of 0.9.

Based on this simulation analysis, we conclude that the sample size is determined by the effect size of age on total alpha power, as this is the smallest effect that was found in the literature. The planned sample size of 200 subjects for the main analysis provides sufficient power for the investigation of both total alpha power and aperiodic signal parameters, even in the case of a considerable proportion of subjects drop out (see Figure 2). This also applies to the validation dataset, which consists of data from 215 subjects.

### 2.11 Secondary analyses

In order to validate the results obtained for the main dataset in the analysis described above, the same analyses will be performed on an openly available dataset [30]. The dataset has been downloaded and no further observations on this data have been made. The dataset consists of EEG recordings of 215 healthy participants comprising a young and an elderly group. In this study, participants were seated in a sound-attenuated Faraday cage, a 16-minute resting-state EEG was recorded (eight blocks of eyes-open condition, eight blocks of eyes-closed condition, each 60s long). EEG was recorded using 62-channel active ActiCAP electrodes, attached according to the international 10-10 system. Data was recorded with a sampling rate of 2500 Hz and referenced to the electrode FCz. Data was bandpass filtered between 0.015 Hz and 1 KHz, and impedances were kept below 5 KΩ.

To ensure this data is comparable to that in the other two datasets, it will be down-sampled to 125 Hz and only the first five of the eight eyes-closed blocks will be used. We will extract data from a timewindow ranging from 2 seconds to 38 seconds after the beginning of each block, as done with the main dataset. Data will be preprocessed, analyzed, and scaled in exactly the same way as described above. Accordingly, the same exclusion criteria will be applied.

For the remaining data, the same statistical procedure will be applied here as described in 2.6. As this dataset is publicly available it has been analyzed several times, yet no other study so far has done similar efforts to investigate EEG age differences in total or aperiodic adjusted alpha power.

### 2.12 Quality & feasibility checks, positive controls & bias minimization

In order to ensure that the obtained results can test the stated hypothesis, we will implement different checks. First, the data quality of the EEG files is automatically and objectively rated by the preprocessing pipeline (see 2.4). Additionally, the grand average topographical distribution of alpha power will be plotted. This also serves as a quality check of the data, as alpha power shows a specific topographical distribution, with maximum power in posterior and occipital electrodes (e.g., refs [6,7]). Furthermore, we also test age differences in total alpha power, which is a prerequisite for testing whether these effects change when adjusting for the aperiodic signal. Replicating previous findings [6,13–17] will ensure that the analyses can also test the additional hypothesis. Pilot analyses (see section 3) serve as feasibility checks for the analysis code, as descriptive support for the principled evaluation of the proposed hypotheses. Positive controls as defined for classical experiments cannot be applied here, as no experimental manipulation is performed in resting state recordings.

All analysis code is prepared using the pilot samples and is shared in an OSF repository (https://osf.io/8e2kd/). The only changes for the analysis of two main datasets, which have not been observed in any way so far, will be done in order to adjust the scripts to correctly handle the different data structure of the two datasets. This will remove all the degrees of freedom for the experimenter.

### 2.13 Anticipated timeline

Data collection of the dataset recorded in our laboratory is ongoing and expected to be completed within the next two months. Preprocessing, analysis and preparation of stage 2 submission including possible exploratory analysis are expected to be finished within 4 months after in principle acceptance.

### 2.14 Data availability plan

All data used for the planned analysis will be stored on the OSF repository https://osf.io/8e2kd/. In this repository, code of the pilot analyses is published. The adapted analyses for the main dataset and the validation dataset will also be publicly available here.

### 2.15 Ethical approval plan

The study was approved by the ethics committee of the canton of Zurich, Switzerland (BASEC-Nr: 2017-00226) and written informed consent was obtained from each subject.

## 3. Pilot data

We conducted a pilot study to test the feasibility of proposed methods, to implement reality checks, to prepare analysis code for the main analyses, and to extract priors for Bayesian analysis in the main study. The pilot study investigated age effects on all the spectral parameters described above in a dataset from a longitudinal cognitive training study [34]. Because of the smaller sample size, the pilot dataset has insufficient power (see Power Analysis section) to conduct robust statistical analyses. Therefore, the pilot data results are presented here exclusively for illustrative purposes and only effect sizes (i.e., Cohen’s d) are reported. We refrain from any interpretation based on these data, which should be based on the two larger independent samples.

### 3.1. Methods

We used a dataset from a previous study published by our group [34] consisting of data from 118 subjects. In this study, participants underwent four weeks of adaptive working memory (WM) training. EEG was recorded twice, once before the onset of the four-week training phase (time point 1) and once after all training was completed (time point 2). EEG recordings included a resting-state condition and a subset of the WM tasks. Only the resting-state EEG data from the first time point was included in this pilot study, as the WM training might otherwise affect the parameters extracted here. The data acquisition, preprocessing, and analysis parameters were identical to those described for the main dataset described in section 2.2, 2.4, 2.5 and 2.7 except that the EEG was recorded with a 256-channel EEG Geodesic Netamps system (Electrical Geodesics, Eugene, Oregon). After the preprocessing, data dimensionality was reduced to the same 70 channels as in the planned study. All parameters described above (see 2.5.1 and 2.5.2) were extracted for each of the 70 electrodes. Of the original 118 subjects, three were excluded from further processing due to poor data quality rating by the preprocessing pipeline (see 2.4).

A final sample size of 115 participant was evaluated (*N*_young_ = 61, mean age = 23.37 years, sd = 3.97 years, age range = 18–35 years, 38 female; *N*_old_ = 54, sd=3.17 years, mean age = 68.42 years, age range = 61–77, 22 female).

#### 3.1.1 Cluster-wise analysis

The same electrode-cluster-based analysis was performed as described in 2.5.3. For this larger electrode cap, matching electrodes for the parieto-occipital cluster were: E101 (Pz), E126 (Oz), E119 (POz), E109 (PO3), E140 (PO4).

To investigate how the age differences in alpha power are affected by the adjustment for the aperiodic signal, the same bootstrapping methods as described above (2.6.1) were applied on the averaged data of the cluster.

In order to receive informative prior distributions for the planned analyses, the same Bayesian models as described above (see 2.6.2) were also fitted to the pilot data.

#### 3.1.2 Electrode-wise analysis

For the additional electrode-wise analysis, each parameter of each electrode was extracted as described above (2.5.1 and 2.5.2, Table 2) and grand-averaged within the two age groups. The results were then plotted as topographical maps to visualize regions of interest for possible differences between the age groups. Additionally, to extract meaningful priors for the main analyses, the brms models were fitted for each electrode. The models were defined as described in equation 1. The models included all extracted parameters as dependent variables (see 2.6.2).

As described in 3.1.1, all parameters were scaled and the same uninformative Cauchy priors were used.

#### 3.1.3 Source level analysis

The same source analysis as described in 2.7 was performed on this dataset.

### 3.2. Results

Due to bad model fits (R^2^ <.90), 6.06% of data points (i.e., only affected data points were removed, not the full subject, as described in 2.5.4) were excluded in the group of old subjects, in the young group, 1.62% of data points were excluded.

In the remaining data, both age groups showed a high model fit over all electrodes for the parameterized power spectrum (*R*^2^_old_ = 0.985, sd = 0.017; *R*^2^_young_ = 0.988, sd = 0.014).

Additionally, in the group of elderly subjects, 1.96% of data points were excluded as they exceeded a threshold of three standard deviations in any of the parameters. In the young group, 1.08% were excluded from further processing.

#### 3.2.1. Cluster-wise analysis

For the parieto-occipital electrode cluster, grand average power spectral components for each age group are visualized in Figure 3:

**Figure 3:**
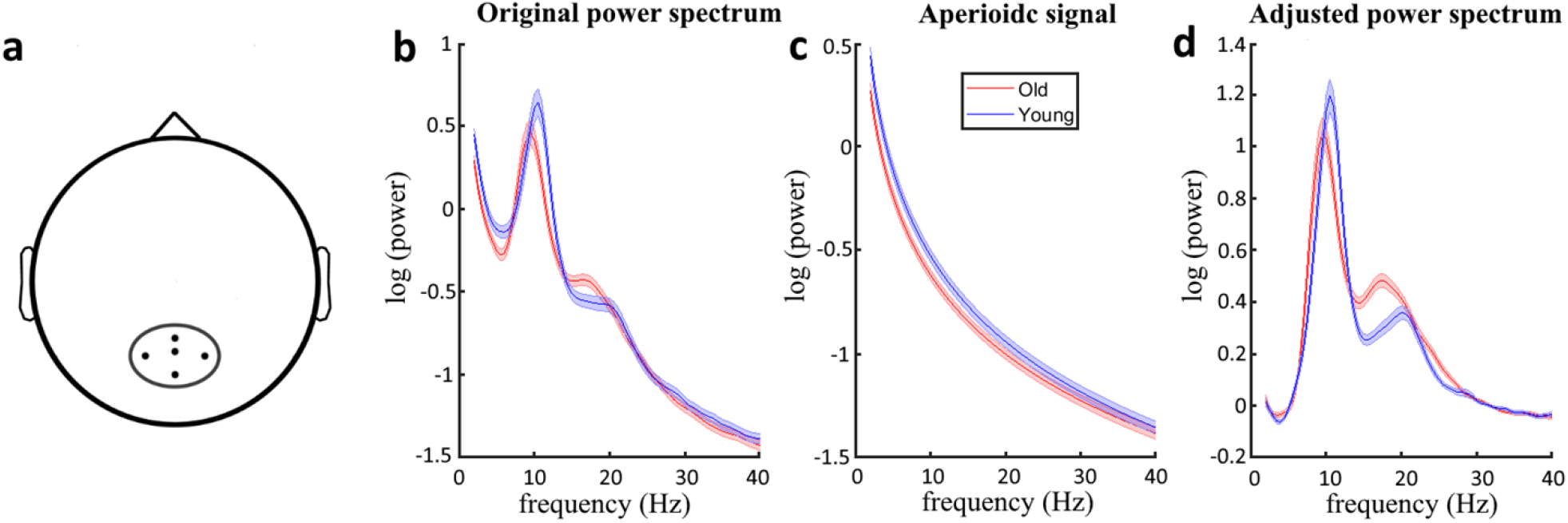
Grand average spectral decompositions in parieto-occipital electrode cluster for both age groups. Shaded areas represent standard error bars. Panel a illustrates the location of the parietooccipital electrode cluster. Panel b shows unadjusted power spectra for each group. Panel c visualizes the fitted aperiodic signal for each age group. Panel d shows aperiodic-adjusted power spectra.

Figure 3 B and C indicate that the decrease of power in age in the range of the alpha oscillation is at least partly driven by decreases in the aperiodic signal. This is also reflected in the bootstrapped age differences in upper and canonical alpha power (Figure 4). While the age differences in the aperiodic adjusted alpha measure are centered at 0 (no age difference), the age difference in the total power spectrum tend to be more negative in this smaller pilot sample.

**Figure 4:**
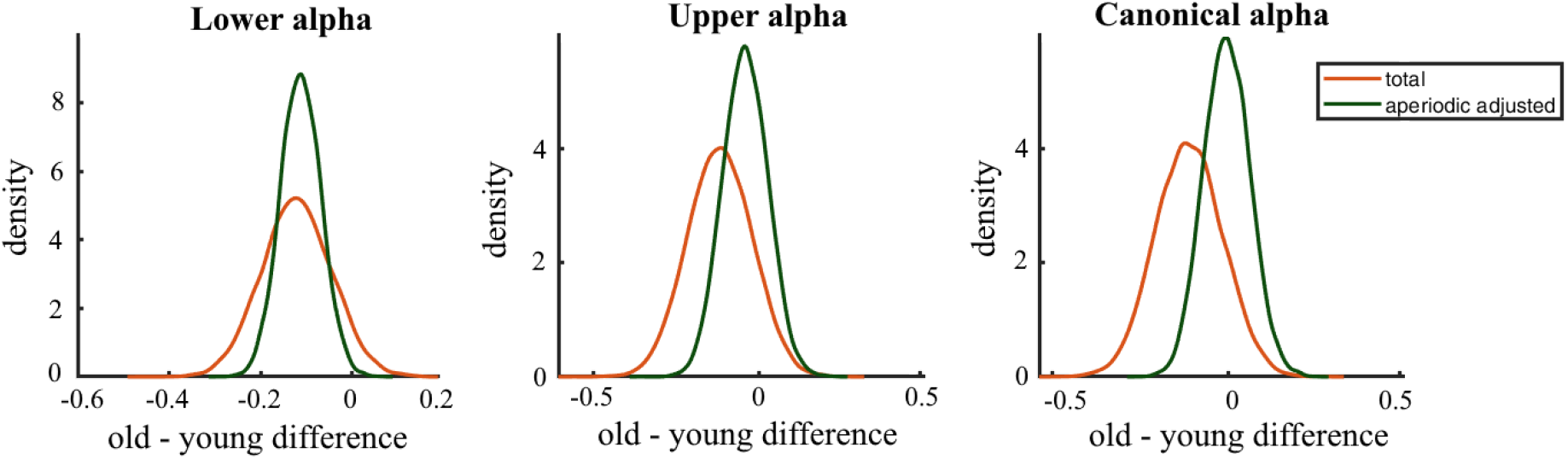
Bootstrapped age difference in lower alpha, upper alpha and canonical alpha band power.

Investigating Cohen’s d for each alpha related measure, decreases in age effects on adjusted power to the corresponding age effect in total power were observed in canonical alpha (d_total_ = 0.23, d_adjusted_ = 0.02) and upper alpha (d_total_ = 0.24, d_adjusted_ = 0.12), while an increase was observed in lower alpha (d_total_ = 0.31, d_adjusted_ = 0.52).

As indicated by Figure 3 C, the aperiodic intercept and slope tend to show a decrease in age in the bootstrapped age differences (see Figure 5). Furthermore, these bootstrapped results indicate a decrease of the IAF in age.

**Figure 5:**
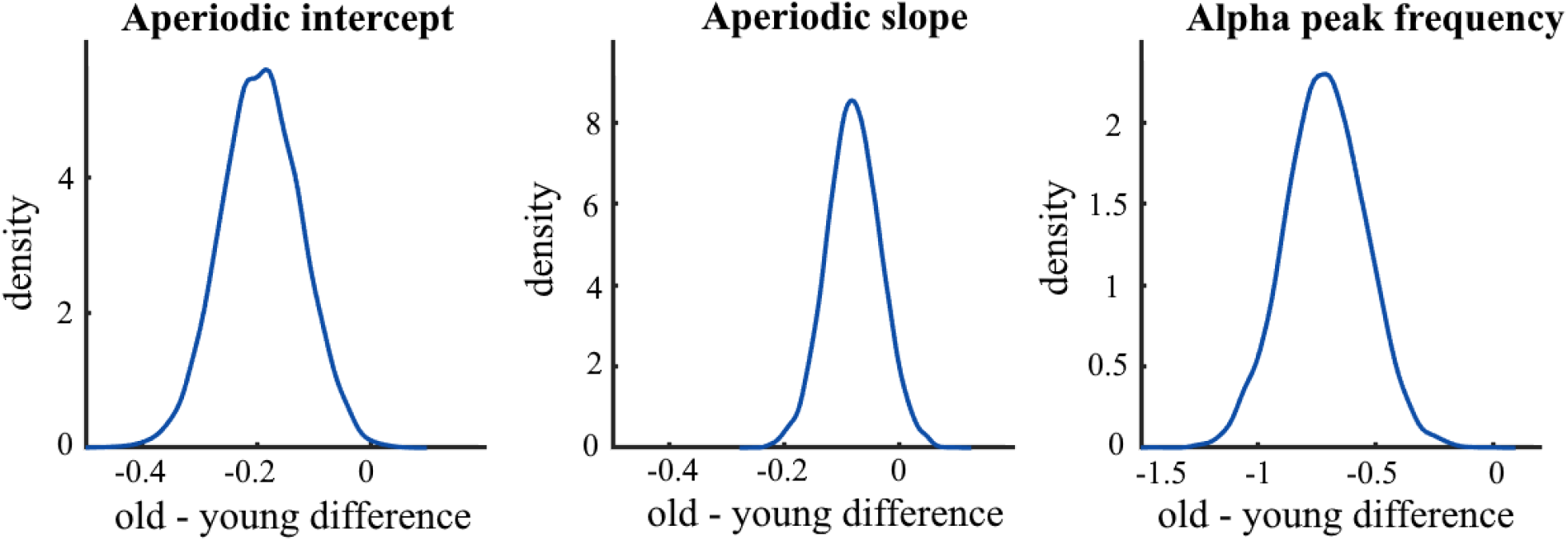
Bootstrapped age difference in aperiodic intercept, aperiodic slope and alpha peak frequency (IAF)

The results of the Bayesian statistical models should not be interpreted as the used sample is not sufficiently powered to investigate age effects in the alpha power measures (see section 2.10). The models were still fitted in order to fix the analyses scripts and to extract priors for the planned analyses. For completeness, the results will briefly be reported in supplementary table 1 (https://osf.io/8e2kd/).

#### 3.2.2 Electrode-wise analysis

Figure 6 shows the age differences (Cohen’s d) for each alpha parameter. The scalp topographies indicate a widespread decrease of the various total alpha power parameters in age. Additionally, when adjusting for the aperiodic signal, an increase of age differences for lower alpha power can be observed, especially in occipital electrodes. In the upper and canonical alpha parameters, the changes in age effects when correcting for the aperiodic signal are more diffuse but tend to decrease in parietal electrodes.

**Figure 6.**
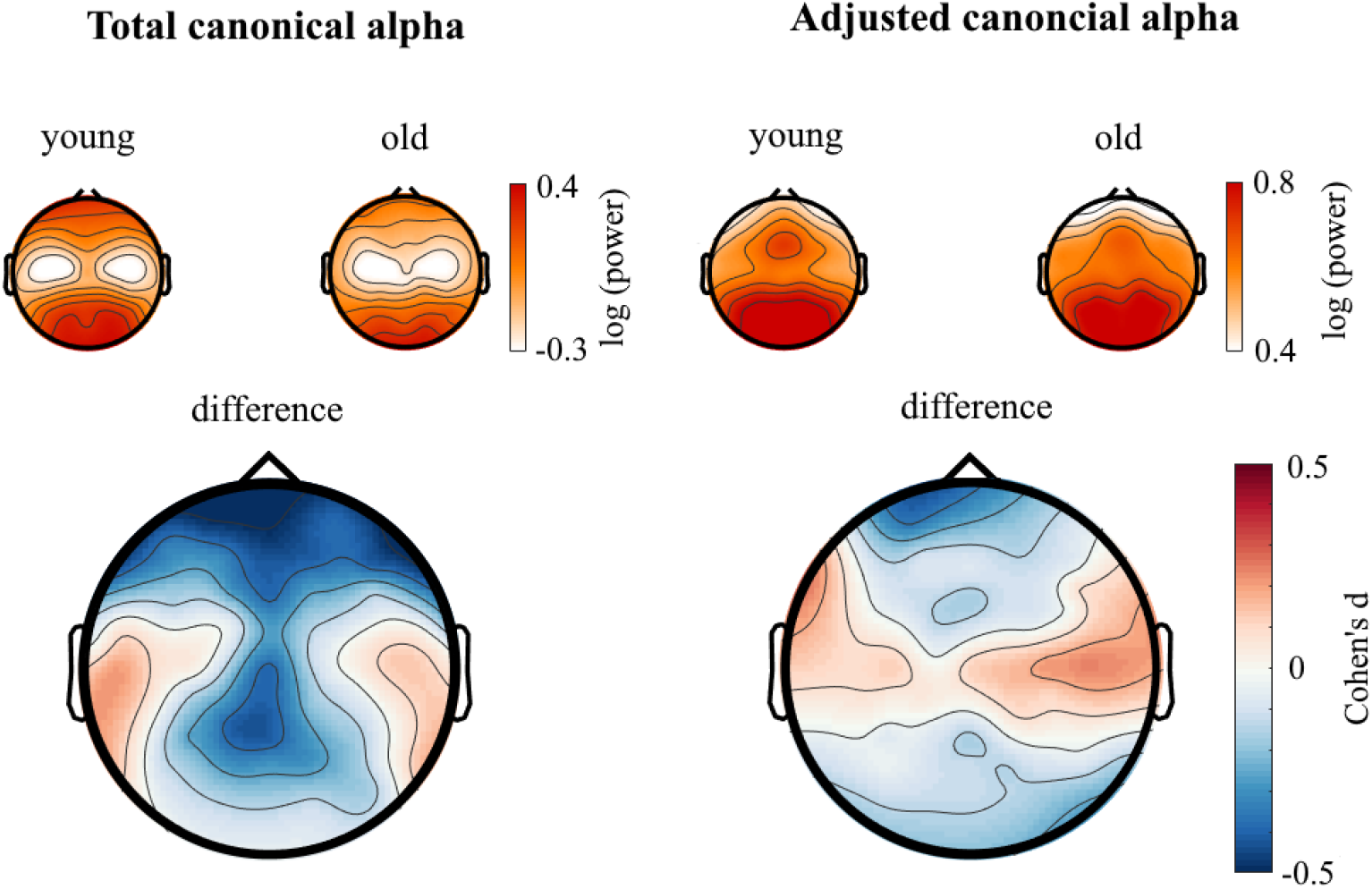
Scalp distribution of age differences (Cohen’s d) for total canonical alpha power and the corresponding aperiodic-adjusted measure. Above each difference plot, topographies of the corresponding measure in log power microvolt are plotted for each age group.

The scalp distribution of the aperiodic signal parameters indicates a widespread decrease in age for both the aperiodic intercept and slope (see Figure 7).

**Figure 7.**
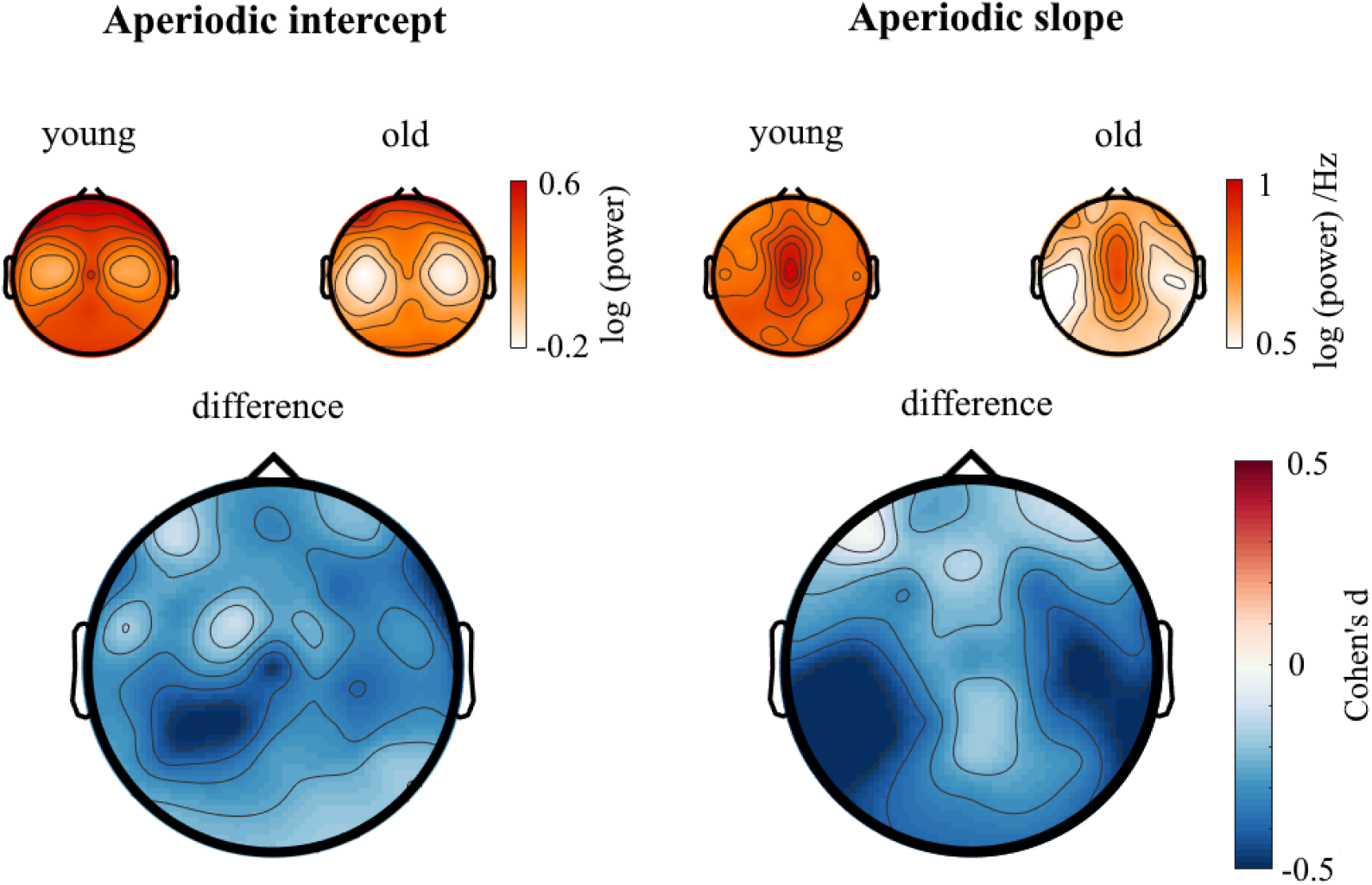
Scalp distribution of age differences (Cohen’s d) of the intercept and slope parameter of the aperiodic signal. Above each difference plot, topographies of the corresponding measure are plotted for each age group.

The topographical plots for lower and upper alpha power age differences and the resulting posterior distributions from the statistical models for each electrode are available in an OSF repository (https://osf.io/8e2kd/). Posteriors are not reported here in detail as they are only used to extract informative priors for the planned main analysis.

#### 3.2.3 Source-level analysis

Source analysis of adjusted canonical alpha power reveals similar spatial distribution as scalp level topographies. Yet, topographical age-related increases in central adjusted alpha power (as indicated in Figure 6) seem to originate from temporal source regions (see Figure 8). These preliminary plots indicate that the previously observed parieto-occipital age-related decreases of alpha power (e.g., refs [13,14]) may no longer be existent in the aperiodic adjusted measure.

**Figure 8.**
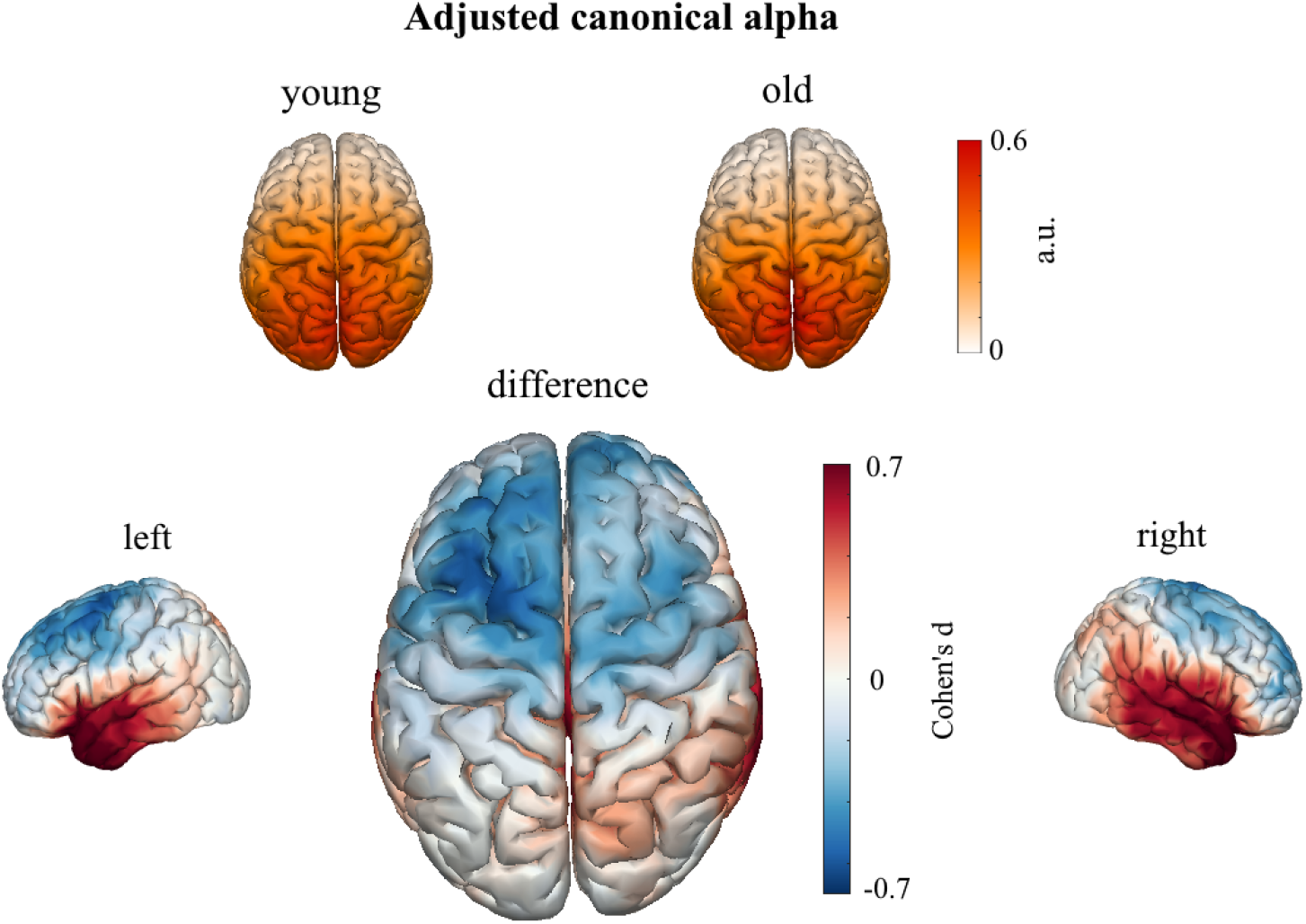
Distribution of age differences (Cohen’s d) for aperiodic-adjusted canonical alpha power in source space. Above the difference plots, spatial distributions are plotted for each age group.

Figure 9 illustrates source reconstructed spatial distribution of the aperiodic slope parameter. Similar to the topographical results (see Figure 7), there are wide spread age differences across the cortex.

**Figure 9.**
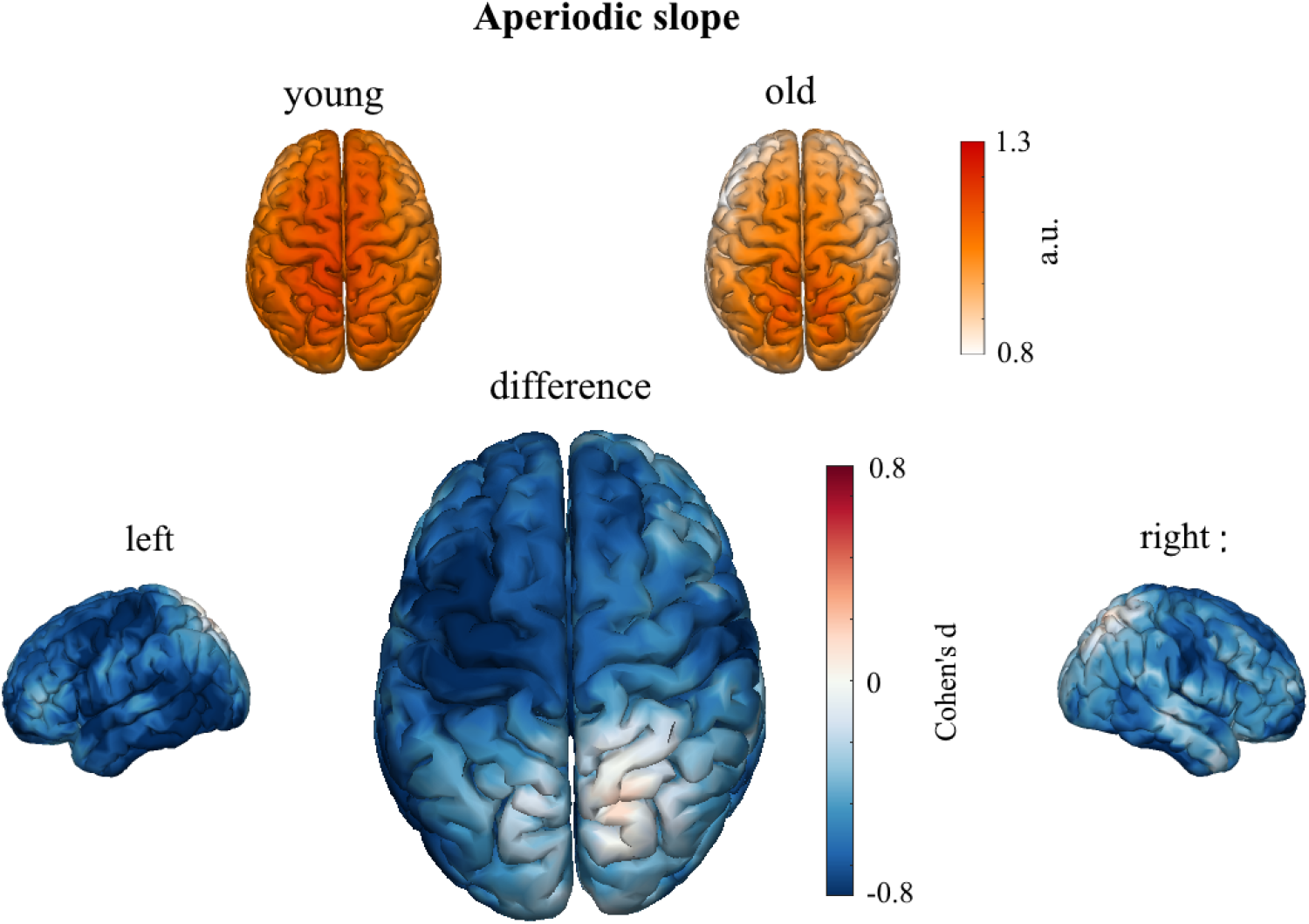
Source reconstructed spatial distribution of the aperiodic slope parameter for each age group (top) and age differences (bottom).

For source reconstructed spatial distribution of lower and upper aperiodic-adjusted alpha power, see supplementary figure 3 and 4 (https://osf.io/8e2kd/).

### 3.3. Evaluation of feasibility of the planned project

Pilot data analysis showed the feasibility of the proposed analysis in the main study, as differences in total alpha power were observed between the two age groups. The statistical models should not be interpreted independently of the main analyses, yet they provide first hints of a replication of previous findings of decreased alpha power in age when analyzing total alpha power in parietal electrodes (e.g., refs [13,14]). Additionally, topographical plots of age differences and the bootstrap statistics indicate that these results may change when adjusting for the aperiodic background signal. At the same time, the analysis of the pilot sample could replicate age differences in the aperiodic signal parameters [23]. This suggests that the aperiodic background signal may introduce a bias in aging studies investigating alpha band power differences, as proposed by ref [19] and awaits further investigation.

